# Cenote-Taker 2 Democratizes Virus Discovery and Sequence Annotation

**DOI:** 10.1101/2020.09.15.298943

**Authors:** Michael J. Tisza, Anna K. Belford, Guillermo Dominguez-Huerta, Benjamin Bolduc, Matthew B. Sullivan, Christopher B. Buck

## Abstract

Viruses, despite their great abundance and significance in biological systems, remain largely mysterious. Indeed, the vast majority of the perhaps hundreds of millions of viral species on the planet remain undiscovered. Additionally, many viruses deposited in central databases like GenBank and RefSeq are littered with genes annotated as “hypothetical protein” or the equivalent. Cenote-Taker2, a virus discovery and annotation tool available on command line and with a graphical user interface with free high-performance computation access, utilizes highly sensitive models of hallmark virus genes to discover familiar or divergent viral sequences from user-input contigs. Additionally, Cenote-Taker2 uses a flexible set of modules to automatically annotate the sequence features of contigs, providing more gene information than comparable tools. The outputs include readable and interactive genome maps, virome summary tables, and files that can be directly submitted to GenBank. We expect Cenote-Taker2 to facilitate virus discovery, annotation, and expansion of the known virome.

## Introduction

Virus hunters have a challenging signal-to-noise problem to consider. For example, animals and bacteria share homologous genes with more amino acid identity than even the most-conserved genes in some virus families (for example, GenBank sequences: polyomavirus Large T antigen [NP_043127 vs. YP_009110677] and 60S ribosomal protein L23 [CUU95522 vs. NP_000969]). Further, there are no universal genes found in all viral genomes that could be used to probe complex datasets for viruses, whereas cellular genomes can be detected through PCR targeting ribosomal genes and alignment of sequences to other single-copy marker genes^1^. Finally, at least hundreds of millions of virus species are likely to exist on Earth^2^, but sequences for only tens of thousands of virus species are deposited in the central GenBank virus database and fewer than 10,000 virus species exist in the authoritative RefSeq database^3^. Sequence space thus covers at, at best, 0.0001% of the virosphere.

Several tools have been developed to detect virus sequences in complex datasets. Strategies include detection of hallmark genes conserved within known virus families (but absent in cellular genomes)^4,5^, detection of short nucleotide sequences believed to be enriched in viruses^6^ (or other machine learning approaches^7,8^), or the ratio of genes common to virus genomes versus genes common to non-viral sequences^9^. Each of these tools has pitfalls that can lead to false positives or false negatives and some tools are limited by minimum sequence length or are only geared to detect a limited range of virus families.

Beyond discovery and detection, *de novo* annotation of contigs representing viruses presents a number of challenges. To list a few, determination of genome topology, accurate calling of open reading frames, determining the virus-chromosome junction in integrated proviruses, resolution of taxonomy, and, especially, accurate annotation of highly divergent homologs of known genes all present technical hurdles^10^. An even deeper problem is the misannotation of some existing GenBank entries. One random example is accession number YP_009506243, which is annotated as a densovirus virion structural protein despite the fact that it is clearly a bidnavirus type B DNA Polymerase. The error has been propagated into more recently deposited bidnavirus sequences (e.g., AWB14612, QJI53745). Relatedly, viral genes and genomes are often misidentified as host sequences^11^. E.g. a mitovirus replication protein (ABK28172) is annotated as an Arabidopsis thaliana protein of “unknown” function.

This manuscript presents version 2.0 of our Cenote-Taker pipeline, which was originally geared toward elementary annotation of viruses with circular DNA genomes^12^. Cenote-Taker2 is a more flexible tool that enables the discovery and annotation of all virus classes with DNA or RNA genomes, starting from genomic, metagenomic, transcriptomic, and metatransciptomic assemblies. It is available for use on Linux terminal and as a graphical user interface (GUI) with free compute cluster usage on <u>CyVerse</u>. The <u>wiki</u> contains a section on suggested parameters for different data types. Cenote-Taker 2 outpaces other currently available annotation tools, providing information for a higher percentage of genes with a higher degree of accuracy, especially for virus hallmark genes, and producing human-editable genome maps that can be opened in any number of genome viewers. Additionally, Cenote-Taker 2 performs better for discovery of viral sequences in complex datasets, with lower false positive and false negative rates than comparable tools.

## Results

### Cenote-Taker2 process overview

A basic run of Cenote-Taker2 requires only a file of contigs from any biological source and a file with metadata that enables submission of annotated sequences to GenBank. A number of optional settings allow users to customize the pipeline. In-depth discussion of the options can be found at the Cenote-Taker2 GitHub <u>repo</u> and <u>wiki</u>. Figure 1 provides a visual of Cenote-Taker2 workflow. First, Cenote-Taker2 analyzes contigs above a user-determined length and detects contigs with inverted or direct terminal repeats. Contigs with terminal direct repeats are circularized and rotated to a position where no open reading frames (ORFs) overlap the wrap-point. An optional step uses raw read data to calculate the average depth of coverage for each contig. All input contigs are then scanned for the presence of a curated set of hallmark genes specific to known virus families. To discover divergent previously unknown viruses in complex datasets, only contigs containing the minimum user-determined number of virus hallmark genes are moved forward for further analysis. For users who have indicated that their input contigs are pre-filtered to only contain viral contigs, all contigs are kept and annotated. Therefore, Cenote-Taker2 can be used simply as an annotation tool, if desired.

**Figure 1:**
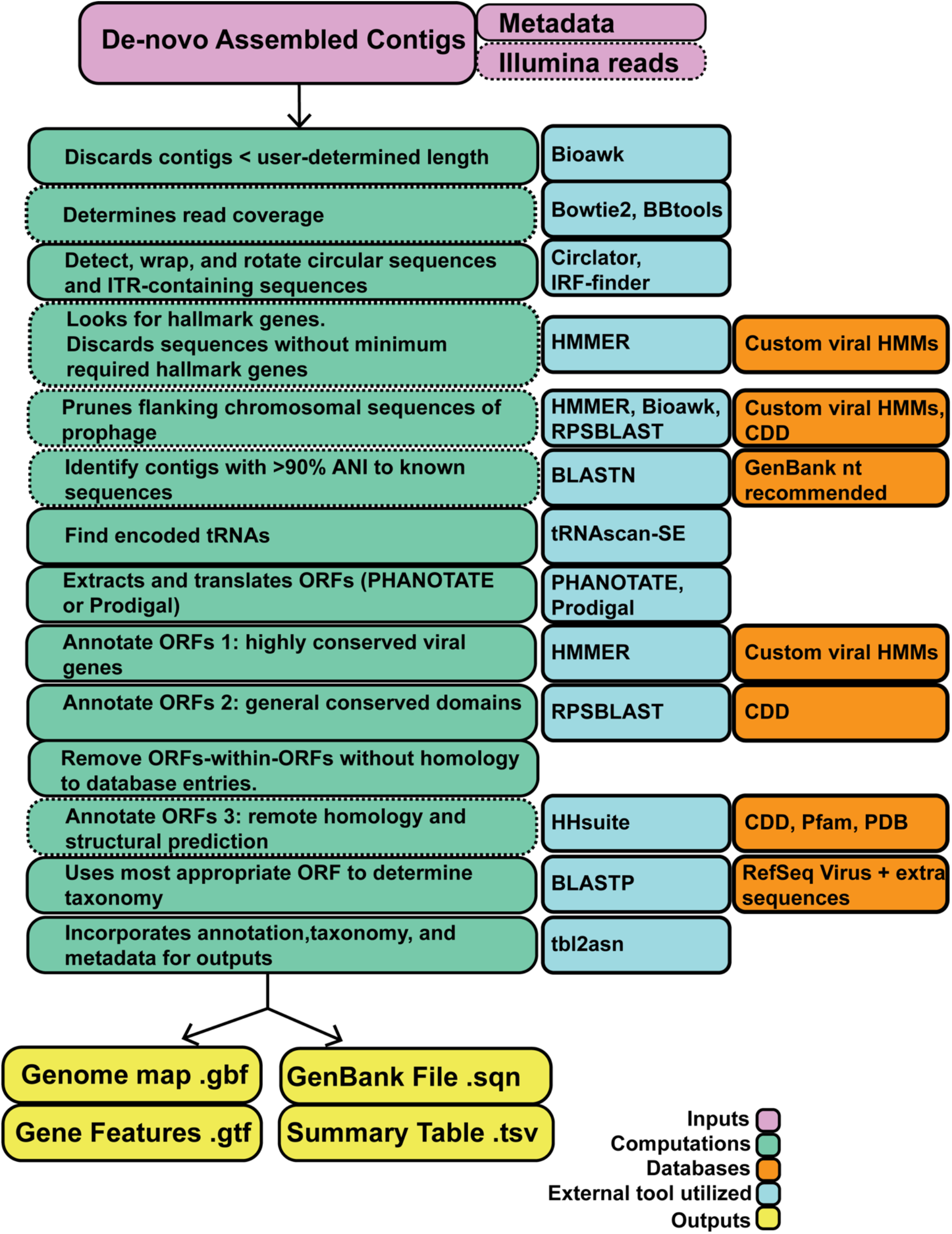
Schematic of Cenote-Taker2 Processes. Visual representation of Cenote-Taker2 virome analysis. Boxes with dashed lines represent optional inputs or processes.

Many viral genomes are integrated into host chromosomes. In datasets likely to containing cellular chromosomes, a single contig might thus contain a virus sequence flanked on one or both sides by a cellular sequence. Users can choose to allow Cenote-Taker2 to prune flanking cellular sequences and generate a genome map for the viral portion of the contig. An optional “known knowns” module conducted at this step queries a nucleotide database, such as GenBank nt, with BLASTN^13^ and marks contigs with at least 90% average nucleotide identity to existing database entries.

Next, candidate tRNA genes are detected and annotated^14^. A tentative taxonomy of each contig is then inferred using BLASTX against a custom database containing Refseq virus and plasmid sequences from GenBank. This taxonomy is used to determine the best ORF-caller (PHANOTATE for putative bacteriophage^15^, Prodigal for other viruses^16^). ORFs are then functionally annotated based on validated datasets using tools for detection of remote homologs (i.e. hmmscan^17^, RPS-BLAST^18^, then HHblits/HHsearch^19^). In these steps, only carefully curated databases (CDD, PFam, PDB, Cenote-Taker2 hallmark database) are queried in order to avoid propagation of mis-annotated sequences in databases such as GenBank nr. For each sequence, a hallmark gene sequence is queried against a reference database of viral proteins using BLASTP. All annotation, taxonomy information, and metadata are combined to generate several outputs. Each contig is represented as an interactive genome map file (.gbf), a gene feature file (.gtf), and a file that can be used for GenBank submission (.sqn). Finally, key information on all annotated contigs is provided in a single virome summary table (.tsv).

### Generation of Hidden Markov Models for Virus Hallmark Genes

Amino acid sequences from public virus databases, RefSeq and GenBank, were downloaded in batches based on family-level taxonomy. Sequences were dereplicated at 70% Identity with CD-HIT^20^, then these representative sequences were clustered using EFI-EST^21^ (pairwise E value cutoff < 1e^−10^). Clusters were visualized in Cytoscape^22^ and multi-lobed clusters were manually divided (removing interstitial sequences) or discarded. Clusters were then further pruned with MCL cluster^23^. Each cluster of three or more proteins was aligned using MAFFT^24^ with default settings. The resulting multiple sequence alignments (MSAs) were used as queries for HHsearch structural prediction and distant homology detection searches against PDB, CDD, and Pfam (80% probability cutoff). MSAs without confident alignment to any models in this search were again used as queries for HHblits against UniProt (80% probability cutoff). Each MSA with a hit in either search was named based on the HHsearch/HHblits top hit and used to generate a hidden Markov model (HMM) using Hmmer. All HMMs were kept for further consideration if the name corresponded to a possible viral hallmark gene (e.g. major capsid protein). All Putative Hallmark HMMs were tested for specificity with a two-step validation by first querying against a negative control database, namely, human proteins from RefSeq, using Hmmer (hmmscan, 1e^−6^ E value cutoff). Second, protein sequences from a variety of human and environmental virome-derived contigs were queried against the database of the remaining HMMs using Hmmer and any proteins with hits to the database were then cross-queried using HHsearch against PDB, CDD, and Pfam. If these proteins had hits to models in these databases that were qualitatively different from the identity of the putative Hallmark HMM, the Hallmark HMM was discarded. To acquire hallmark genes not represented by GenBank or RefSeq database, genomes from the human gut virome database^25^ and virome assemblies from seawater^26^ were translated, and amino acid sequences were processed as above. Finally, HMMs from pVOGs^27^ and PFAM^28^ were considered and validated in the same manner. Some replication-related Hallmark HMMs were later removed because they were similar to genes typically found on plasmids or conjugative transposons. Virion structural, virion processing, and virion packaging gene HMMs are used by Cenote-Taker2 with a cutoff of 1e^−8^ and genome replication gene HMMs are used with a cutoff of 1e^−15^.

### Cross-comparison of Currently Available Virus Annotation Modules

At present, VIGA^29^ is the only publicly available genome annotation tool specifically designed for viruses. To compare VIGA’s genome annotation function to Cenote Taker 2, we arbitrarily chose four “challenging” viral genomes as case studies (Fig. 2). For the a newly described amoeba-tropic virus called Yaravirus^30^ (Fig. 2A), only Cenote-Taker2 could discern an annotation for any genes, with the major capsid protein (MCP), packaging ATPase, and replicative helicase all being recognizable. For a crAss-like phage assembled from a human gut metagenome dataset (Fig. 2B), Cenote-Taker2 again annotates more genes than VIGA. For a soil-associated levivirus^31^, only Cenote-Taker2 could identify the capsid/maturase gene and the RNA-Dependent RNA Polymerase (RDRP). In this case, the levivirus RDRP gene lacked a stop codon and this prevented VIGA, but not Cenote-Taker2, from calling the ORF correctly. For a fish-associated inovirus, only Cenote-Taker2 was able to identify the packaging ATPase (ZOT), MCP, and Attachment protein. For the most important functional annotations for each genome, supporting evidence is shown from HHpred and DELTA-BLAST (Fig. S1). Direct genome map outputs from VIGA and Cenote-Taker2 are available for each genome (Supplemental Files 1-8).

**Figure 2:**
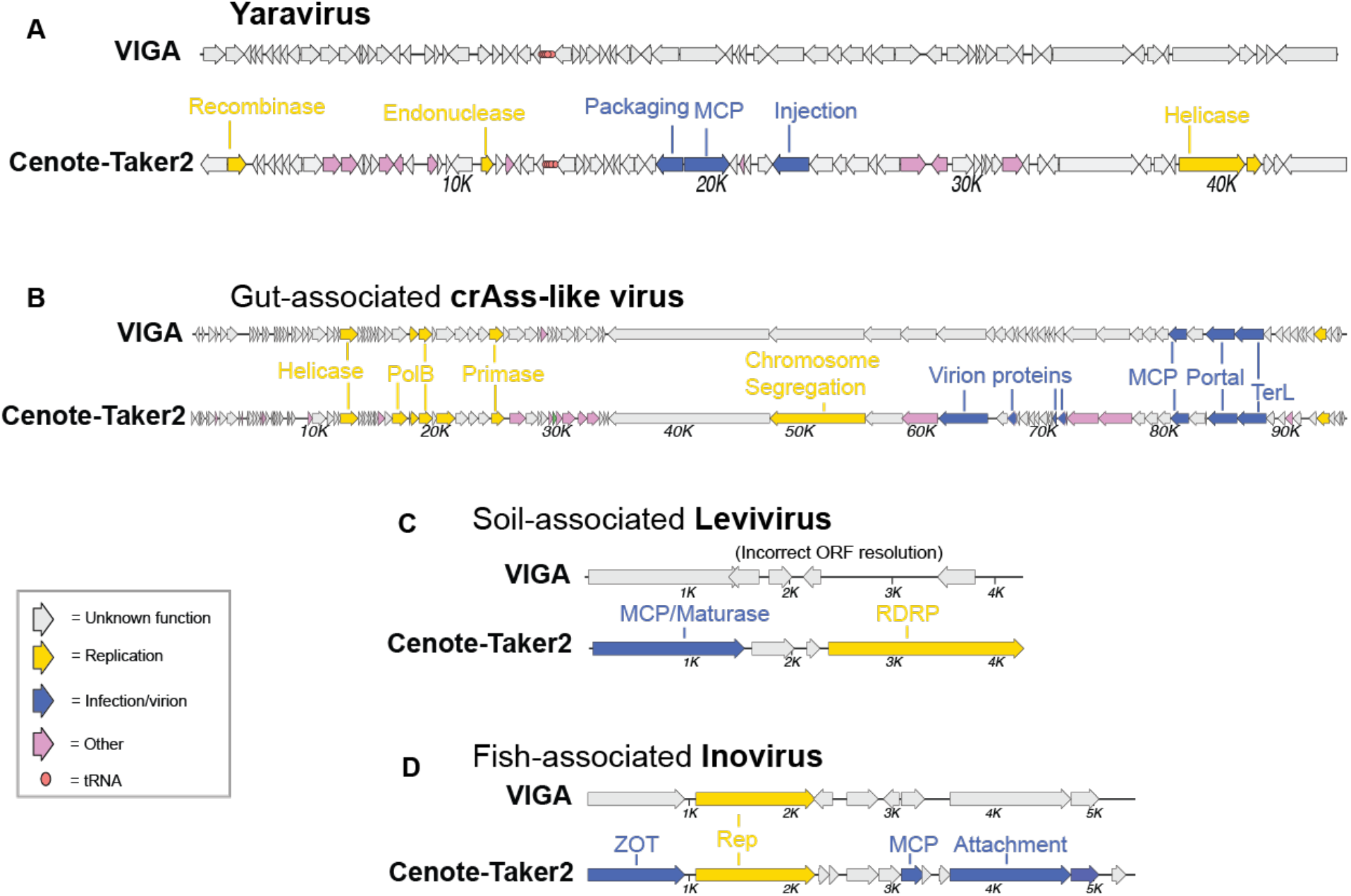
Comparison of genome maps from VIGA and Cenote-Taker2. Cenote-Taker2 and VIGA were run with optimized options (see methods). (A) Yaravirus (accession MT293574) is a newly reported midsize DNA virus found in amoebae. (B) crAss-like viruses are tailed phages. The species shown here was assembled from a human gut metagenome SRA dataset SRR6128032. (C) Leviviridae sp. isolate H1_Bulk_28_FD_scaffold_59 (accession MN033558) is a levivirus genome identified in a soil metatranscriptome. (D) Inoviridae sp. isolate ctba29 (accession MH616818) is an inovirus found in a haddock virome dataset.

### Comparison of Virus Discovery Module

Cenote-Taker2 was compared to three leading virus discovery tools, each with its own method for detecting viral sequences. Like Cenote-Taker2, VirSorter^4^ uses a virus hallmark gene detection approach. One limitation is that it is only designed to detect bacteriophages. DeepVirFinder^6^ uses a machine learning approach to find short nucleotide motifs common in viral sequences. An additional pipeline, Non-Targeted^9^ (used for “Uncovering Earth’s Virome”^32^), compares predicted protein sequences encoded by a contig to a curated set of known viral and cellular proteins. A limitation of Non-Targeted is that it only considers contigs greater than 5 kb, while the other tools have no strict minimum length. The main categories of complex datasets that might be searched for new viruses are: assembled contigs derived from DNA samples enriched for viral sequences (DNA virome), RNA samples enriched for viral sequences (RNA virome), DNA from unenriched samples (genomes and metagenomes), or RNA from unenriched samples (transcriptomes and metatranscriptomes). Additional parameters to consider are the fact that ssDNA viruses may require a second strand synthesis step for DNA samples, and multiple displacement amplification (MDA) may selectively enrich for circular viral genomes through rolling circle amplification (RCA) effects^33^. Examples of each category of dataset were assembled and scaffolded (see methods), and contigs greater than 1000 nucleotides were analyzed with the four virus discovery pipelines. Cenote-Taker2 outperformed all other discovery tools for finding contigs with genes encoding for virion components (i.e. “structural”) or replication genes for each type of dataset (Fig. 3, Fig. 4).

**Figure 3:**
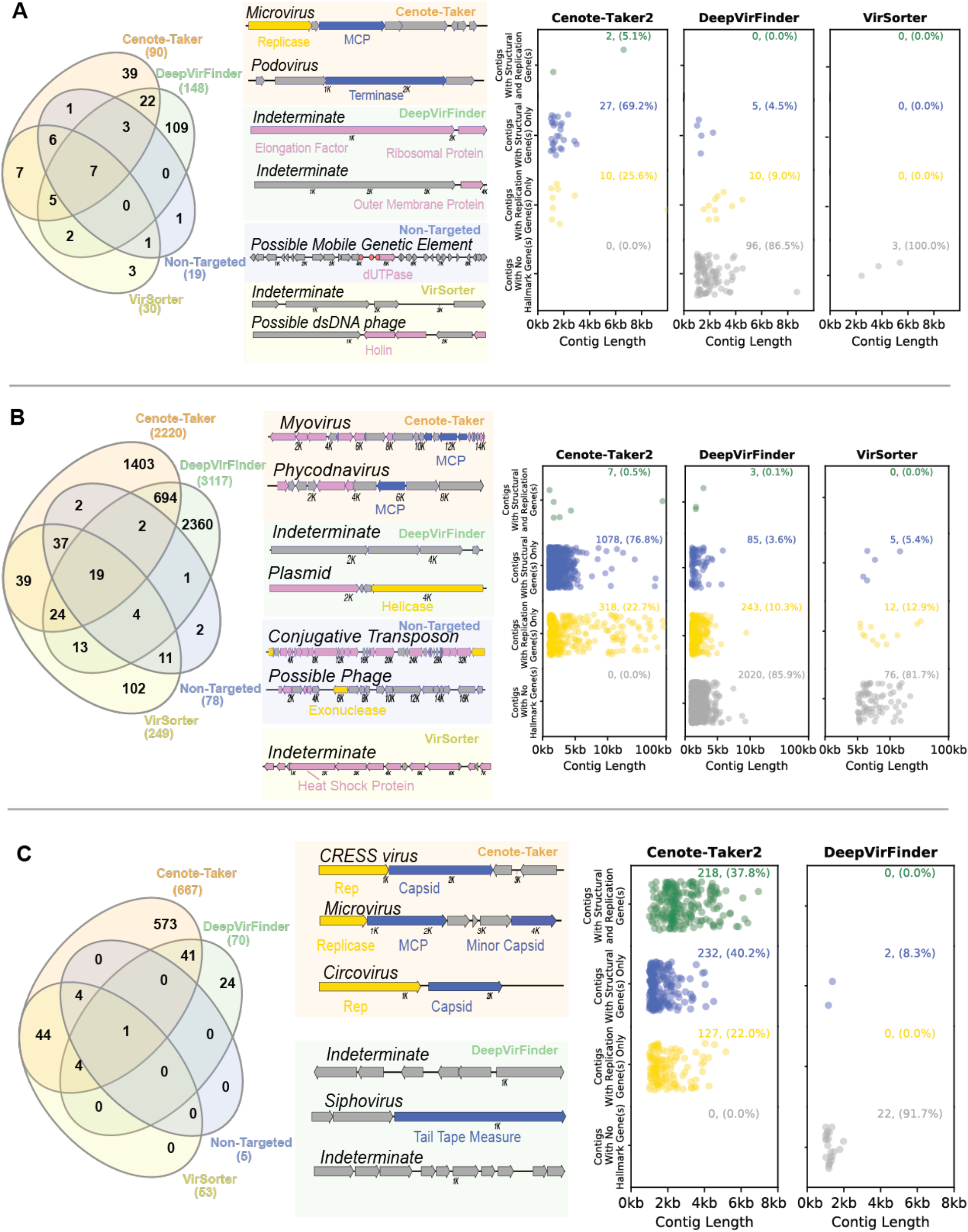
Comparison of virus discovery tools for DNA datasets. Contigs > 1000 nucleotides were analyzed using four virus detection/discovery pipelines. (A) A dataset for human stool enriched for nuclease-resistant DNA in virus-sized particles (SRR6128021). Left panel: Venn diagram displaying the overlap of contigs identified as viruses by the various pipelines. Middle panel: maps showing representative examples of unique calls from each pipeline. Left panel: scatter plots display the length distribution for calls unique to each software package. Rows of plots show contigs that contain both virion structural genes and DNA or RNA replication-associated genes, only one of the two categories of gene, or neither. (B) Similar display of a dataset for Amazon River water samples analyzed with shotgun total DNA sequencing (ERR2338392). (C) Similar display of a dataset for wastewater enriched for nuclease-resistant DNA in virus-sized particles amplified using MDA/RCA (SRR3580070).

**Figure 4:**
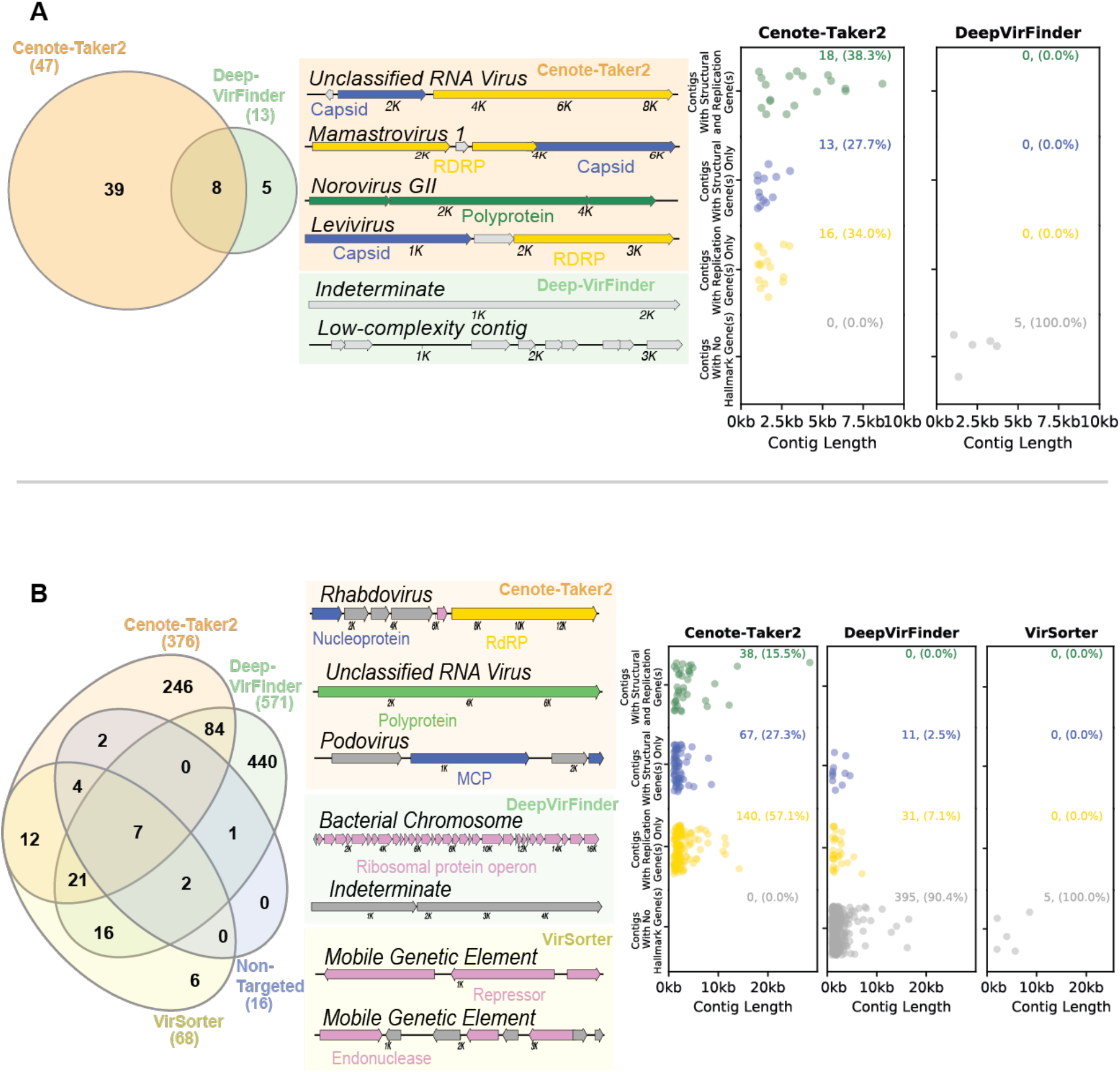
Comparison of virus discovery tools for RNA datasets. Contigs > 1000 nucleotides were analyzed using four virus detection/discovery pipelines. (A) A dataset for sewage analyzed with viral particle capture and RNA sequencing (ERR3201762). Left panel: only Cenote-Taker2 and DeepVirFinder had any virus calls. Middle panel: maps of representative examples of contigs which each pipeline uniquely called as viral. Right panel: Contig attribute chart showing only contigs called uniquely by Cenote-Taker2 and DeepVirFinder. (B) A dataset for stool from a Tasmanian Devil with whole metatranscriptome RNA sequencing (SRR8048119). Left panel: comparison of the overlap of contigs the various pipelines designated as viral. Middle panel: maps of representative examples of contigs which each pipeline uniquely called as viral. Right panel: contig attribute chart showing only contigs called uniquely by Cenote-Taker2, DeepVirFinder and VirSorter.

For a dataset of a DNA virome dataset (i.e., virus-like particle enrichment with nuclease digestion, followed by DNA sequencing) from human gut^34^, DeepVirFinder had the most total virus calls (149), of which 109 were unique to DeepVirFinder (Fig. 3A). However, when unique calls were analyzed in more detail by functionally annotating genes with RPS-BLAST and HHpred, most contigs uniquely called by DeepVirFinder were found to lack any virus hallmark genes, making these calls potential false positives. Cenote-Taker2 also had many unique calls (39) and all uniquely called contigs encoded a virion structural/packaging or replicative viral hallmark gene (Fig. 3A), implying that Cenote-Taker2 has higher specificity for viruses. A similar trend can be seen for a large metagenome (dsDNA) assembly from Amazon river water^35^ (Fig 3B). For the metagenome dataset DeepVirFinder again had the most calls, but most of the unique calls were ambiguous upon closer inspection.

A different pattern was seen for a waste water sample from which DNA was subjected to second strand synthesis through MDA/RCA^36^ (Fig. 3C). For the amplified DNA dataset, Cenote-Taker2 detected more total calls (5.2X all other tools combined) and more unique calls (23.8X all other tools combined), all of which have at least one type of hallmark gene. Single-stranded DNA viruses are highly abundant members of many microbial communities^12,37,38^ making Cenote-Taker2’s discovery module a particular advance for researchers interested in these viruses.

Cenote-Taker2 also detected more RNA viruses from an sewage RNA virome dataset (virus-like particle enrichment with RNA sequencing) than DeepVirFinder (Fig. 4A). Furthermore, nearly all the unique calls from DeepVirFinder are low complexity sequences or only contain unrecognizable ORFs. VirSorter and Non-Targeted did not detect any viruses in this dataset, consistent with the fact that they were not designed to detect RNA viruses.

Metatranscriptome samples are perhaps the most complex category of dataset because they are expected to contain RNA virus genomes, transcripts from DNA viruses and other mobile genetic elements. For a metatranscriptome dataset for a Tasmanian devil stool samples^39^, DeepVirFinder called the most viruses, but over 90% of its unique calls were found to indeterminate or clear false positives. Cenote-Taker2 detected both RNA and DNA viruses in this set (Fig. 4B). Overall, Cenote-Taker2 appears to show superior sensitivity and specificity for bona fide virus contigs in all types of dataset.

### Prophage Pruning Module

When the Cenote-Taker2 prophage pruning module is selected, linear contigs are assigned ORF calls via Prodigal, then ORF are iteratively searched with 1) HMMSCAN of the custom virus hallmark gene database, 2) HMMSCAN of the custom common virus gene database, and 3) RPS-BLAST of CDD. Each gene is then considered to be 1) a virus hallmark gene, 2) a common viral gene (hit in the custom “common” (but not hallmark) virus gene database or hit in CDD of a domain found in 10 or more RefSeq Caudovirales genomes or hit in CDD with ‘PHA0’ prefix), 3) a common host gene (all other CDD hits), or 4) an unknown gene (no hits in any of these databases). Based on the coordinates of the ORFs and their categorization, each nucleotide position in the contig is scored as likely virus or likely host. Bases within virus hallmark or common viral genes are scored as 10. Bases within unknown genes, which are more common in viruses, are scored as 5. Bases in intergenic regions are scored as 0 and bases within known bacterial genes are scored as −3. The sum of 5 kb windows tiled every 50 bases is calculated, then scores are smoothed based on the scores of adjacent windows. Contig segments with one or more consecutive windows with a positive score are resolved, and segments containing virus hallmark genes are designated as viruses or virus fragments.

To show that Cenote-Taker2 can identify virus genomes from order Caudovirales, which commonly occur as prophages and encode a variable range of accessory genes, all 3493 putatively complete Caudovirales genomes were downloaded from virus RefSeq. Each sequence was fed to Cenote-Taker2 with the prophage pruning module on. First, 3487/3493 genomes were identified as having at least one viral hallmark gene in the Cenote-Taker2 database (Fig 5A), with almost all also having several hallmark genes. Three of the six hallmark-negative sequences (NC_042064, NC_042059, NC_042564) were incomplete genome fragments. One (NC_002670) was a phage satellite^40^. One (NC_023591) was a mobile genetic element with many conjugative genes but no genes annotated as virion structural or packaging genes, suggesting that it is non-viral. The last (NC_029050) was a sequence that had almost no callable ORFs and is perhaps a degraded prophage relic.

**Figure 5:**
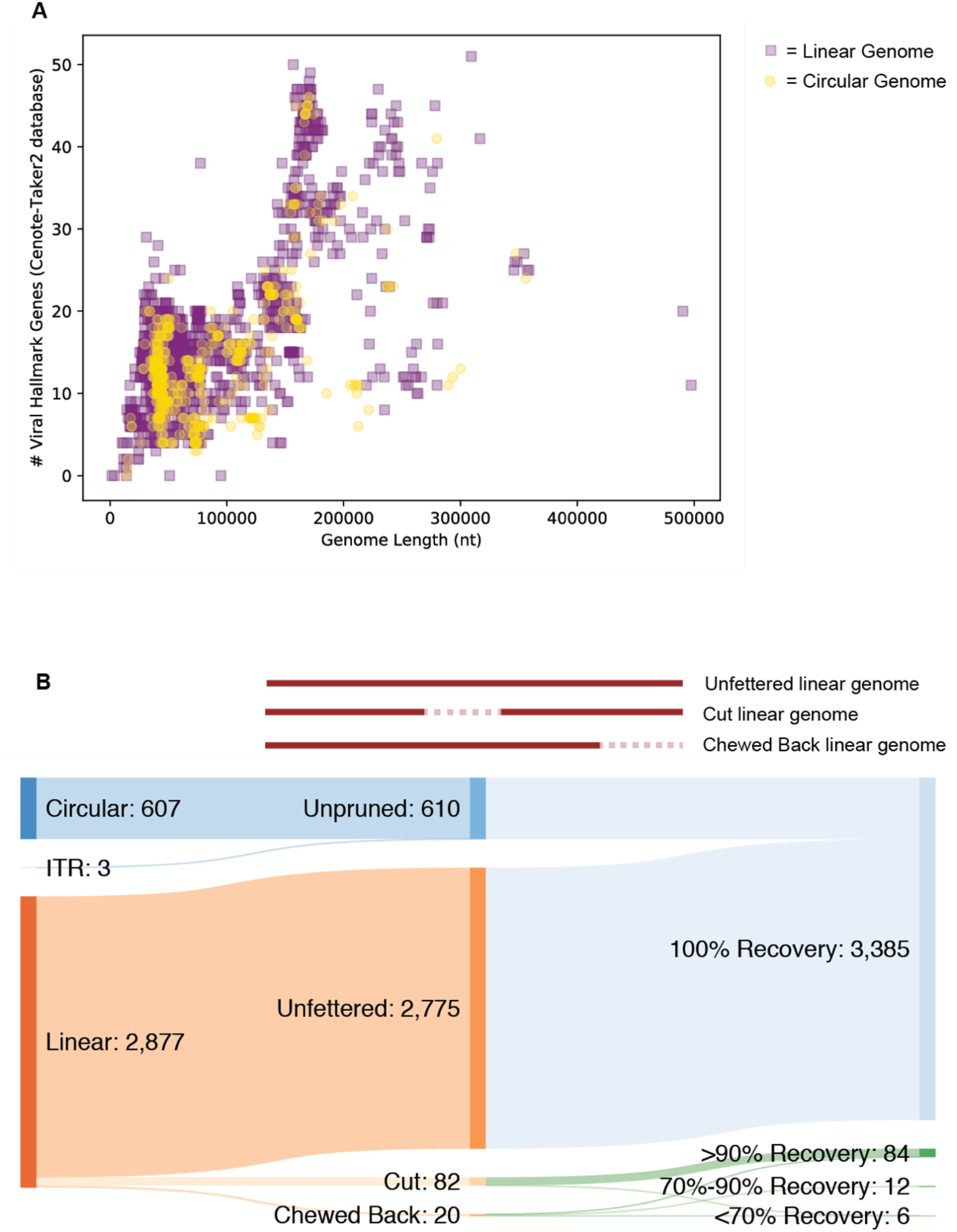
Cenote-Taker2 detects and does not prune the vast majority of Caudovirales genomes. (A) 3493 putatively complete Caudovirales genomes were downloaded from NCBI RefSeq. Each genome is represented as a dot with its length on the x-axis and the number of genes called as viral hallmarks by Cenote-Taker2 on the y-axis. (B) Top: schematic visualizing perturbations/lack thereof to genomes caused by Cenote-Taker2 pruning module. Bottom: Sankey diagram of input/output of 3487 Caudovirales RefSeq genomes.

To investigate whether the Cenote-Taker2 pruning module might truncate Caudovirales sequences, the length of each genome was analyzed before and after pruning (Fig. 5B). Of the 2877 genomes eligible for pruning (610 of the 3487 were recognized as circular or flanked by ITRs and not eligible), 96.5% (2775/2877) were kept intact by the pruning module. 2.9% (82/2877) of genomes were “cut” in the middle because the pruning module removed loci falsely determined to be nonviral (each case only had one cut region). Over 90% of the original genome was kept after pruning in all but one cut genome. 0.7% (20/2877) of genomes were “chewed back” from one end. Seven of 20 had >90% recovery, twelve of 20 had 70%-90% recovery, and five of twenty had <70% recovery.

As a more applicable example, the main chromosome of a Bacteroides xylanisolvens genome (genome assembly ASM654696v1) was analyzed with prophage pruning on. Prophage calls and virus genome maps are shown in Figure 6, with three apparently full-length siphoviruses and one full-length microvirus prophage being discovered.

**Figure 6:**
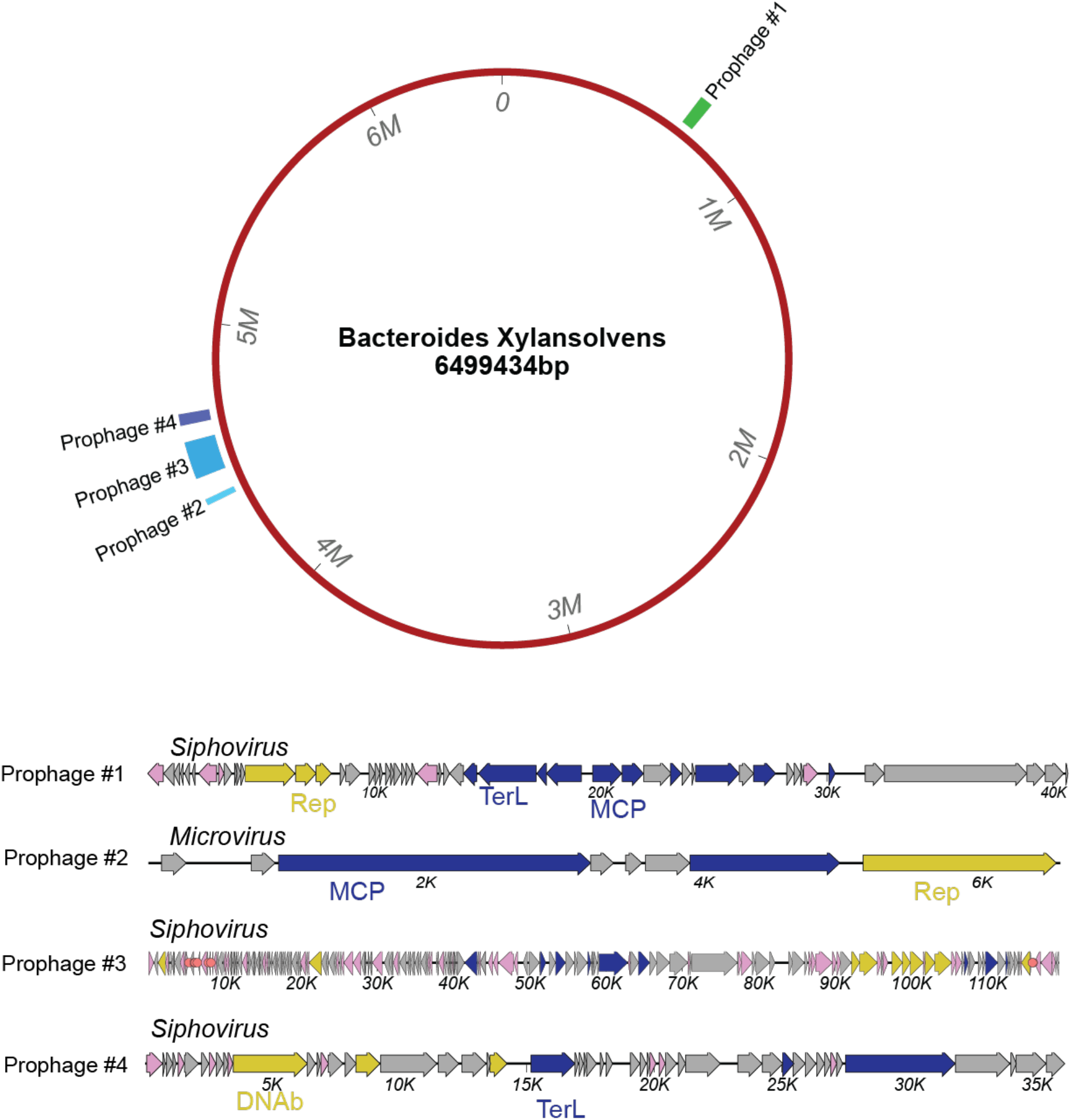
Cenote-Taker2 analysis of *Bacteroides xylanisolvens* genome (GenBank assembly: ASM654696v1) with prophage pruning. The circular map represents the *B. xylanisolvens* genome annotated with coordinates of the prophage called with Cenote-Taker2. The Cenote-Taker2-generated map of each prophage is shown.

## Discussion

We expect Cenote-Taker2 will prove useful to scientists who wish to detect and annotate viruses, including divergent previously unknown virus species, in large and complex datasets. Cenote-Taker2 empowers users both with the ability to easily discover viruses in complex datasets as well as the ability to quickly analyze candidate viruses through visualization of annotated genome maps in any available genome or plasmid map viewer. Further, combining discovery and annotation should dovetail nicely with other techniques to cluster viral sequences into species level^41,42^ or higher taxonomic levels^10,43^, especially when conducting pairwise comparisons of virus genomes within or between taxa^44^. Furthermore, because Cenote-Taker2 eases submission of annotated genomes to GenBank, even those who don’t use Cenote-Taker2 will indirectly benefit by having a larger, better-annotated, central sequence database.

Two known annotation challenges of viral coding regions that are not resolved with Cenote-Taker2 are ribosomal frameshifting, which is documented in some RNA viruses and dsDNA bacteriophage, and intron-containing genes, which are common in eukaryotic viruses. Other non-canonical translated features could be missed, as well. We are not aware of current tools for automating the resolution of these features. Additionally, functional annotation is somewhat limited by only using databases with well-curated gene families, which precludes the annotation of newly characterized gene families not yet in gold standard databases. However, the custom database of over 3000 HMMs of viral hallmark genes developed with Cenote-Taker2 goes beyond PFAM, PDB, and CDD databases and mitigates some of these limitations.

Cenote-Taker2 outperforms other currently available virus discovery pipelines for a variety of reasons. While both VirSorter and Non-Targeted employ hidden Markov models of viral genes to some extent, it’s likely that the models developed for Cenote-Taker2 represent more of the diversity of viral hallmark genes. Further, since contigs are penalized by Non-Targeted if they contain common chromosomal genes, contigs representing a virus sequence flanked by a chromosomal sequence might be discarded instead of pruned. DeepVirFinder uses a fundamentally different approach, looking for nucleotide k-mers of different lengths to determine if a contig is a virus. Two reasons why this approach can fall short are: (1) nucleotide sequence space may be unable to adequately capture the vast diversity of virus genomes (2) DeepVirFinder was trained on “virome” assemblies. Physical enrichment of viruslike particles is notoriously difficult^45^, so some training datasets may have been contaminated with host sequences. Moreover, it is known that some sequences, even in very clean virus-like particle preparations, are not viruses but mobile genetic elements that parasitize viral capsid machinery^46^. Two new tools capable of virus discovery have come out very recently, only after we completed our analysis^47,48^.

While there are likely new “types” of yet-to-be discovered viruses encoding novel capsid and replication genes, Cenote-Taker2 can readily be updated to include new hallmark gene models. For example, a new model was made for the highly derived RNA-dependent RNA polymerase gene of the proposed new family *Quenyaviridae*^49^.

## Methods

### Cenote-Taker2 Code

Cenote-Taker2 was written in Bash, Perl and Python. All scripts can be accessed on <u>GitHub</u>. In-depth discussion of use-cases and considerations can be found on the <u>Wiki</u>. Installation uses Conda to manage packages^50^. BLAST and Hmmer databases developed for this tool can be found on <u>Zenodo</u>.

### Annotations of Challenging Viral Genomes

Cenote-Taker2 was fed these genomes with default settings except “--enforce_start_codon False” was used. Since VIGA default settings are highly stringent, several custom options were used to improve annotation: “--diamondevalue 1e-04 --diamondidthr 25 --hmmeridthr 25 --blastidthr 25”. Genome maps were visualized with MacVector 16.

### Virus Discovery Comparison

Reads from each sequencing run were trimmed with Fastp^51^, assembled with Megahit^52^ (default settings), and scaffolded with SOAPdenovo2^53^. Cenote-Taker2 hallmark gene HMM (Hmmer) database (updated April 21st, 2020) was used with viral hits having one or more detected hallmark genes. The Cenote-Taker2 script requires a p-value of 1-e08 as a minimum threshold for virion structural genes and 1e-15 for replication genes. VirSorter was used with “virome” settings and categories 1, 2, 4, and 5 were kept. DeepVirFinder was used with the default training set and p value threshold of 0.005. Non-Targeted Pipeline was used with default settings. Comparisons were run on April 23rd, 2020.

Contigs uniquely called by Cenote-Taker2 were determined to either have hits in the virion structural or viral genome-packaging gene HMM set and/or in the virus genome replication-associated gene HMM set. Putative viral contigs called uniquely by other sources were annotated with Cenote-Taker, using RPS-BLAST with the CDD database (1e^−4^ e value cutoff) and HHsearch (80% probability cutoff) with CDD, PFam, and PDB. All annotated genes were scanned for names of viral replication or virion structural genes and domains.

## Acknowledgements

The authors would like to acknowledge early users of Cenote-Taker2, especially Leen Beller, Kema Malki, and Keir Balla, for their many helpful suggestions.

## Data availability

GenBank submissions of “third party annotations” of virus genomes require a DOI number of the associated manuscript. The accession numbers of annotated genomes from this manuscript will therefore be provided when this manuscript gets a DOI number. In the meantime, see supplemental .gbf files.

---

**Supplementary Figure 1:**
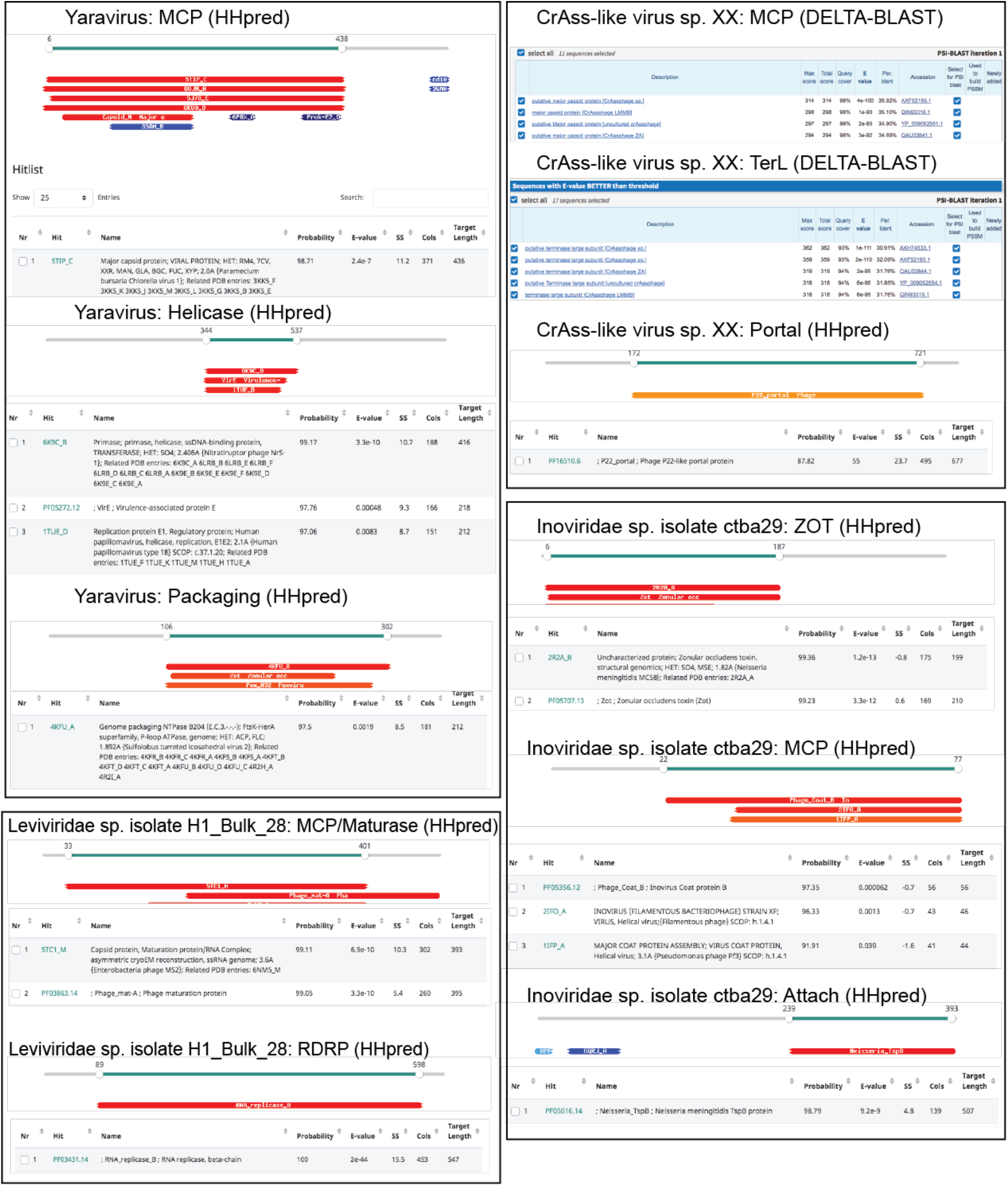
Supporting evidence for functional annotations made by Cenote-Taker2. From genomes displayed in Figure 2, key genes that were annotated by Cenote-Taker2 were queried with HHpred^54^ or DELTA-BLAST to check validity of the functional annotations.

